# Effect of beneficial sweeps and background selection on genetic diversity in changing environments

**DOI:** 10.1101/2022.08.29.505661

**Authors:** Sachin Kaushik

## Abstract

Neutral theory predicts that the genetic diversity within a population is proportional to the census population size. In contrast, observed genetic diversity for various species is much lower than theoretical prediction (Lewontin’s paradox). The selective sweeps and background selection, reduce the genetic variation at the linked neutral sites and have been studied considering the environment to be selectively constant. However, in a natural population, the selective environment varies with time. Here, we investigate the impact of selective sweeps and background selection on neutral genetic diversity when the selection coefficient changes periodically over time. The reduction in genetic variation due to selective sweeps is known to depend on the conditional fixation time. Here, we find that the effect of changing environment on conditional mean fixation time is most substantial for the randomly mating population than the inbreeding population with arbitrary inbreeding coefficient. We also study the effect of background selection on neutral sites when the selection coefficient of linked deleterious mutation change periodically in time. In the slowly changing environment, we find that neutral heterozygosity is significantly different, and the site frequency spectrum has a different shape than that in the static environment.

## 1 Introduction

There has been much interest in understanding how the neutral genetic diversity patterns are shaped for a long time. It has been observed that pairwise diversity in various natural species is several orders of magnitude lower than predicted by the neutral theory (Kimura, 1971) -known as Lewontin’s paradox (Lewontin, 1974). The indirect effects of selection on the linked neutral variation are one possible contributor to resolving Lewontin’s paradox (Buffalo, 2021; Charlesworth and Jensen, 2022).

When a beneficial mutation spreads throughout an entire population, the linked neutral variants can hitchhike along with it; this is referred to as selective sweep (Maynard Smith and Haigh, 1974; Berry, 1991; Stephan *et al*., 1992; Stephan, 2019). Other variants not linked to the selected mutation are subsequently lost from the population, reducing genetic diversity at the linked neutral sites. The genetic diversity can also be reduced due to the elimination of the recurrent deleterious mutations from the population, known as background selection (Charlesworth, 1993) because the deleterious mutations are selected against and subsequently eliminated from the population, which also eliminates the linked neutral variants. Several analytical and simulation studies have been conducted to investigate the effect of background selection (Charlesworth, 1993; Hudson and kaplan, 1995; Gordo *et al*., 2002) and selective sweeps (Braverman *et al*., 1995; Stephan, 2019) on the linked neutral genetic diversity considering environment to be selectively constant.

Natural environments, on the other hand, are not static and change over time. It is therefore important to ask how the genetic diversity at the linked neutral sites is affected by selective sweeps and background selection in the changing environment. We primarily want to know how the levels of genetic variation change in the selectively changing environment. As selective sweeps and background selection effects are strong when recombination is low, we work in low recombination regimes throughout the article.

In a recent study (Kaushik and Jain, 2021), the effect of a continually changing environment on the genetic diversity in a randomly mating population was investigated using a two-locus model with one neutral and one selected locus. It was discovered that the Maruyama-Kimura symmetry (Maruyama, 1974; Maruyama and Kimura, 1974) for conditional mean fixation time in static environment does not hold in a changing environment, and the deleterious sweeps are strongly affected due to varying selection. The beneficial sweeps, on the other hand, are robust for slow or moderate changes in the selective environment (Kaushik and Jain, 2021). Here, we extend these results for selective sweeps to a more general case of inbreeding population. We use diffusion theory to calculate the conditional mean fixation time for the inbreeding population. When the inbreeding coefficient is high, the deleterious and beneficial sweeps are weakly affected in the changing environment, and the changing environment effects are most significant in a randomly mating population.

Because the deleterious mutations are more likely to be lost than fix in population, we investigate the impact of background selection on the shape of the site frequency spectrum (SFS) in changing selective environment. In a finite asexual population, fittest class of individuals is eventually lost from the population (Muller, 1964; Felsenstein, 1974). Previous research has shown that in slow Muller’s ratchet regime and static environment, the reduction in the genetic diversity caused by strong selection against the recurrent deleterious mutations is equivalent to the neutral population with a reduced effective population size (Charlesworth, 1993; Hudson and Kaplan, 1995) in the absence of recombination. However, the effect of background selection in the population where Muller’s ratchet clicks fast is much less studied (Gordo *et al*., 2002). In this article, we will investigate the effect of background selection for arbitrary selection strength in slowly changing environment. We give simple analytical expression for change in heterozygosity when Muller’s ratchet clicks slowly and study the fast Muller’s ratchet regime numerically.

We find that, even in a slowly changing environment, genetic diversity at linked neutral sites differs significantly from that in a static environment when the selected site contains deleterious mutations; in particular when the population fluctuates between zero and a negative selection coefficient, the shape of the time-averaged SFS is found to differ from the shape of the SFS with a time-averaged selection coefficient.

## 2 Selective sweeps in changing environments

The reduction in genetic variation caused by selective sweeps is significant in regions with low recombination and depends on the time it takes for the mutation to fix in the population (Maynard Smith and Haigh, 1974; Barton, 2000). In this section, we discuss the conditional mean fixation time, which is calculated by averaging the time taken by the mutation to fix into the population only in the trajectories destined to fix, when the selective environment is time-dependent and a single locus is under selection.

### 2.1 Diffusion theory

When the environment is selectively constant, the conditional mean fixation time for beneficial mutation with selective advantage +*s* and dominance coefficient *h* is identical to that of deleterious mutation with selective disadvantage −*s* and dominance 1 − *h* (Maruyama, 1974; Maruyama and Kimura, 1974). In the static environment, this symmetry also holds for the inbreeding population (Glémin, 2012). It has recently been demonstrated that this symmetry breaks for randomly mating populations in a changing environment (Kaushik and Jain, 2021). In this section, we consider the case when inbreeding occurs in the population.

Consider a diploid population of constant population size *N*. The biallelic locus experiences time-dependent selection coefficient, 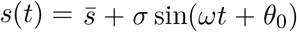 that varies periodically with time, where 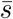 is the time-averaged selection coefficient, 0 *< θ*_0_ *<* 2*π* is the initial phase, *σ >* 0 is the amplitude of the oscillation, and *ω* is the rate of change of environment. The three genotypes, *AA, Aa* and *aa* have frequencies *p*^2^ + *fpq*, 2*pq*(1 − *f*) and *q*^2^ + *fpq*, and fitnesses 1, 1 + *hs*(*t*), and 1 + *s*(*t*) respectively (Charlesworth, 2010; Crow and Kimura, 2009) where *h* is the dominance coefficient (0 *< h <* 1), and *f* is the inbreeding coefficient (0 *< f <* 1). The *p* and *q* are the frequency of allele *A* and *a*, respectively.

The backward Fokker-Planck equation for the randomly mating population (Eqn. A4 in Kaushik and Jain (2021)) for mean fixation time for a mutation with initial frequency *x* at time *t*_0_, can be generalized for inbreeding population, which satisfies

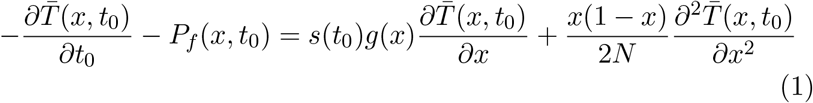

where *g*(*x*) = (*x* + *h*(1 − 2*x*) + *f* (1 − *h* − *x* (1 − 2*h*))), and *P*_*f*_ is the eventual fixation probability.

The above equation does not appear to be exactly solvable, so we use first-order perturbation theory for the approximate solution. Previous research (Kaushik and Jain, 2021) indicates that the effects of a changing environment are significant when the mutation is on-average neutral, so we focus on-average neutral mutation. In the slowly changing environment, *ω* ≪ *N*^*−*1^, *s*(0), for on-average neutral mutation 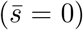 the mean fixation time and eventual fixation probability are approximated as 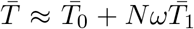, and *P*_*f*_ ≈ *P*_0_ + *NωP*_1_. Here, 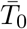 and *P*_0_ are the mean fixation time and fixation probability in the static environment; and 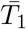 and *P*_1_ are the leading order deviation in *ω* in the mean fixation time and fixation probability, respectively. These are calculated in Appendix A.

The conditional mean fixation times for the slowly changing environment are shown in Fig. 1, where the data is obtained by numerically solving (A.3), (A.9), and (A.10). We find that the Maruyama-Kimura symmetry does not hold for the inbreeding population, and deleterious mutations are more affected than the beneficial ones by changing environment. Thus, in the case of an inbreeding population, the qualitative behavior of conditional mean fixation time is similar to that of a randomly mating population in a slowly changing environment.

**Figure 1:**
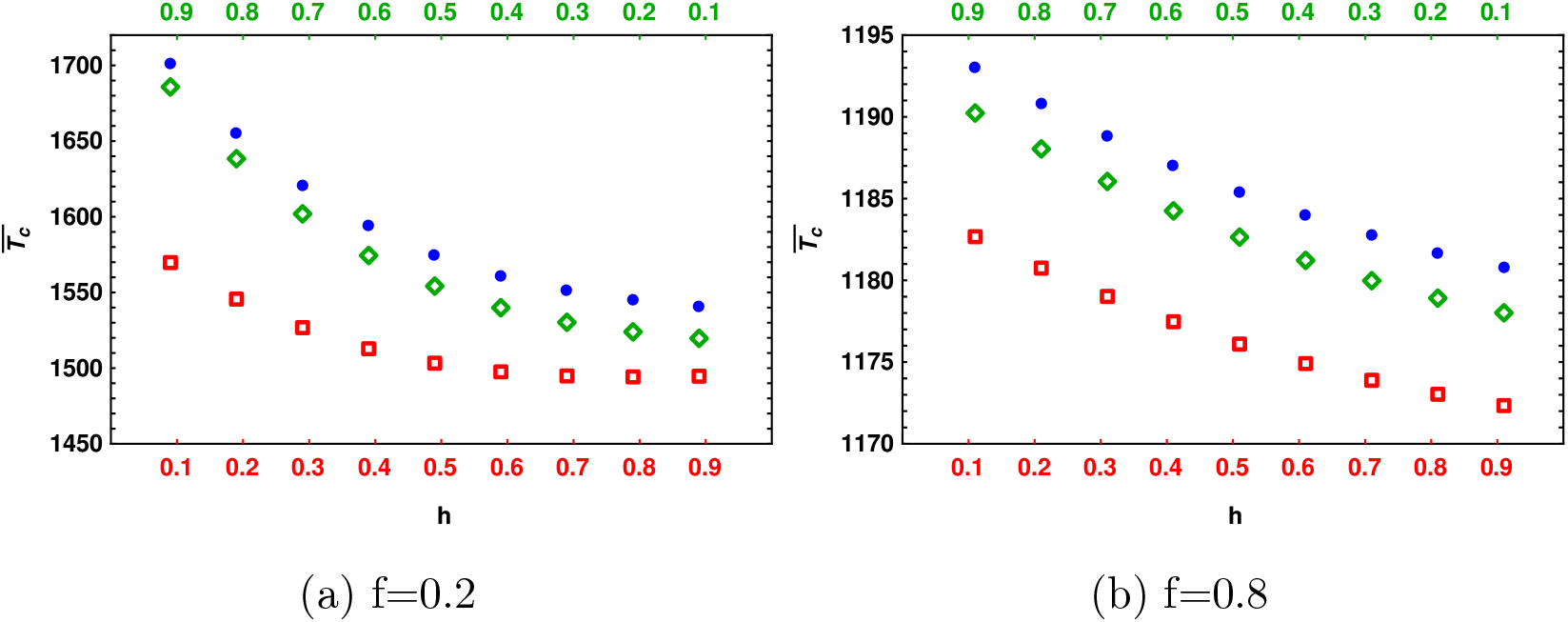
The variation of conditional mean fixation time 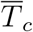 with dominance parameter (*h*) for moderate selection in static 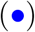 and slowly changing environment where selection coefficient is on-average neutral 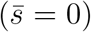, and given by *s*(*t*) = *σ* sin(*ωt* + *θ*) (diamonds) and −*s*(*t*) (squares) respectively. The other parameters are *N* = 2 × 10^3^, *Nω* = 0.05, *σ* = 0.01, *θ* = *π*/4, (a) *f* = 0.2 and (b) *f* = 0.8. When inbreeding is high, the conditional mean fixation time for both beneficial and deleterious is mildly affected in the slowly changing environment.

Whereas, unlike the randomly mating population where the recessive deleterious mutations are strongly impacted, deleterious mutations, whether recessive or dominant, are mildly affected due to the slowly changing environment when the inbreeding coefficient is high, as shown in Fig. 1b. This can be understood qualitatively using the following argument. In a randomly mating population, the recessive deleterious mutations are strongly affected because they have a higher fixation probability than dominant mutations (Haldane, 1927) allowing recessive mutations to have a less strong tendency to fix quickly as compared to the dominant deleterious mutation, and former mutations can continue to segregate in the population for long periods of time, making them more sensitive to the changing environment (Kaushik and Jain, 2021). However, when the inbreeding coefficient is high, the fixation probability for the deleterious mutations is weakly dependent on the dominance coefficient (see A.7, A.8). The trajectories for recessive, as well as dominant deleterious mutations, are then exposed to the changing environment for nearly the same amount of time and thus affected in the similar manner. Thus, these results suggest that in the case of a highly inbred population in a slowly changing environment, the deleterious and beneficial sweeps produce similar genetic diversity patterns.

### 2.2 Semi-deterministic theory: strongly beneficial mutations

The solution of diffusion equation does not have a closed form in the presence of selection and thus cannot be used to calculate the fixation time distribution. However, we can use semi-deterministic theory for the beneficial mutations when the population size is large (2*Ns* ≫ 1), as recently used by Martin and Lambert (2015), and Kaushik and Jain (2021). In a static environment, the conditional mean fixation time, 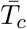, for the mutation with selection coefficient +*s* and dominance coefficient *h* is the same for the mutation with selection coefficient +*s*, and dominance 1 − *h* for arbitrary inbreeding coefficient. In other words, 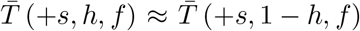 (Ewing *et al*., 2011; Glémin, 2012). This symmetry is preserved in the changing environment for the panmictic population in the strong selection limit, as recently demonstrated by Kaushik and Jain (2021). Here, we will explore the *h* to 1 − *h* symmetry when inbreeding occurs.

The allele frequency distribution, Φ_f_(*p, t*|*p*_0_, 0), that the mutation frequency is *p* at time *t*, given that the initial mutation frequency is *p*_0_ at time *t*_0_ obeys the following forward Kolmogorov equation (Risken, 1996),

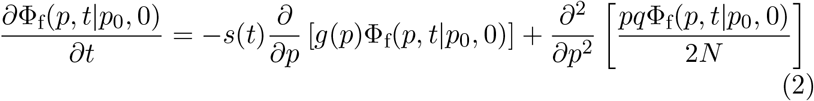

where, *g*(*p*) = *s*_*a*_(1 − *p*) + *s*_*b*_*p, s*_*a*_ = *h* + (1 − *h*)*f* and *s*_*b*_ = 1 − (1 − *f*)*h*, respectively. For *p* → 0, the (2) reduces to a Feller diffusion equation (Feller, 1951a) which can be solved exactly. Conditional on fixation, Φ_f_(*p, t*|*p*_0_, 0) converges to a stationary distribution at large times, which gives the eventual fixation for the beneficial mutation having frequency *p*_0_ at time *t* = 0 as

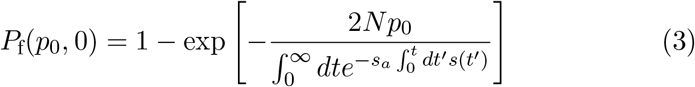

#### Slowly changing environment

In a slowly changing environment, the fixation probability for an on-average neutral mutation 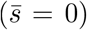, when the mutation occurs in the positive cycle of the selection coefficient with initial phase, *θ*_0_ = *ωt*_0_ (0 *< θ*_0_ *< π*), is obtained by a Taylor series expansion of (3) to the linear order in *ω*. The fixation probability is then given as

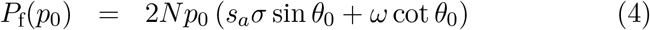

The deviation in the fixation probability is independent of the dominance parameter and inbreeding coefficient. As previously discussed, the conditional mean fixation time is closely related to the eventual fixation probability; thus (4) suggests that a slowly changing environment is expected to affect conditional mean fixation time mildly.

The conditional mean fixation time is approximated as 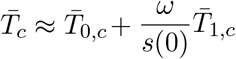 when the environment changes slowly. For a single mutation, and keeping terms up to the order 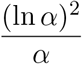, the first order correction to the conditional mean fixation time is given by (see Appendix B for details),

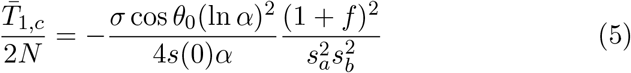

For a randomly mating population, *f* = 0, the above expression reduces to Eqn. (20) in Kaushik and Jain (2021). In slowly changing environment, the leading order term in 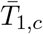 has *h* → 1 − *h* symmetry. In other words, like in randomly mating population, 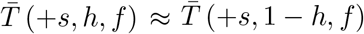 holds in the slowly changing environment for the inbreeding population also. The deviation in the conditional mean fixation time, 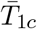, due to changing environment is maximum when *f* = 0, as depicted in Fig. 2.

**Figure 2:**
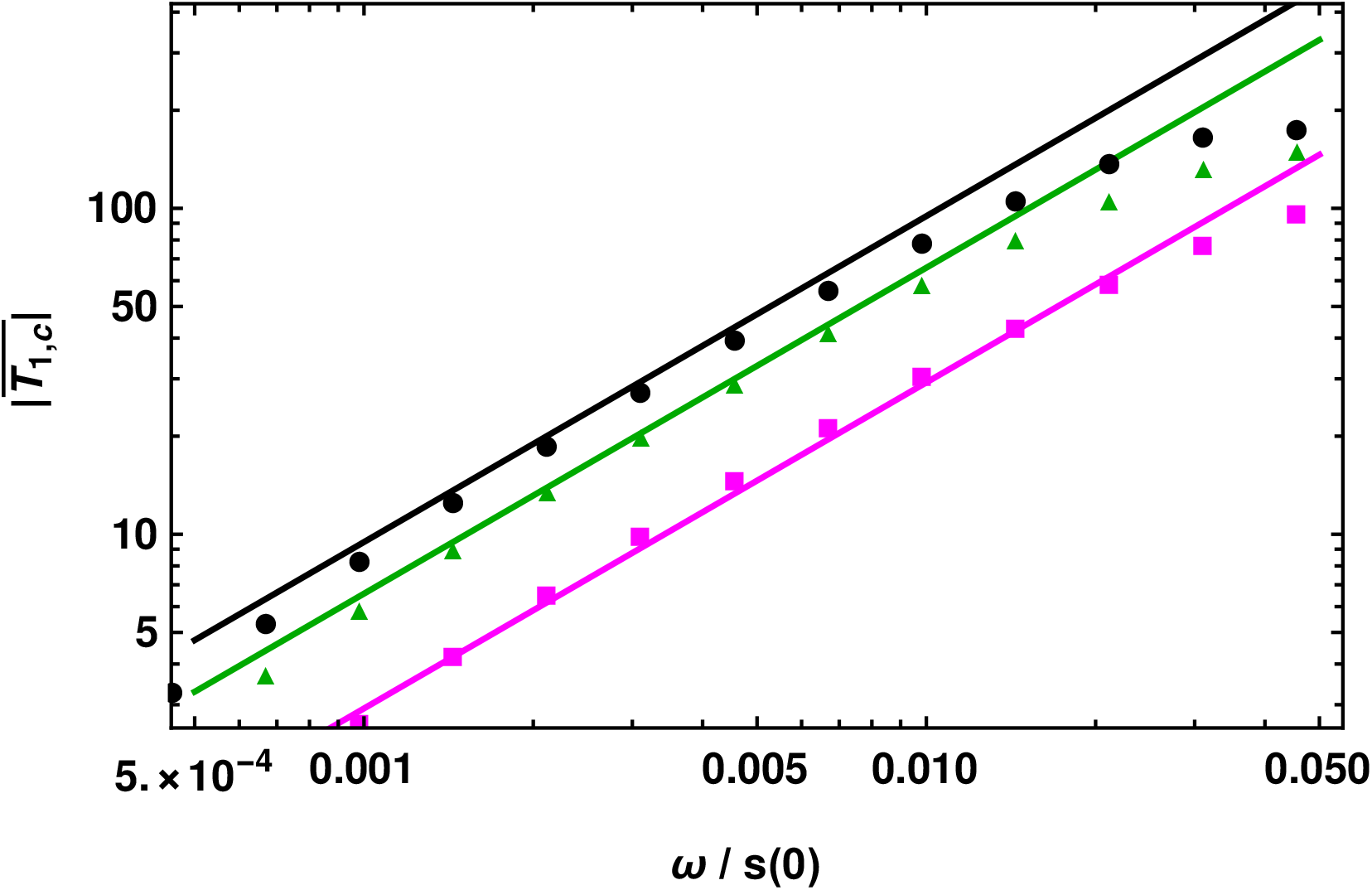
Strongly beneficial mutations: The deviation in the conditional mean fixation time, 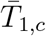 in the slowly changing environment with selection coefficient 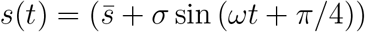 when the mutation is beneficial at all times for *f* = 0 (black), *f* = 0.2 (green), and *f* = 0.8 (magenta) to show that the deviation is maximum for randomly mating population. The points are obtained by numerically solving (B.1). The other parameters are *N* = 10^5^, 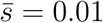, *σ* = 0.007 and *h* = 0.5 respectively.

Fig. 2 shows that the conditional fixation times are affected very mildly in the wide range of environmental chaange. Thus, the results of the beneficial sweep in the changing environment for a randomly mating population are robust in the inbreeding population.

## 3 Background selection in changing environment

In the previous section, we have shown that the effect of changing environment on deleterious sweeps is strongest for the randomly mating population. In this section, we therefore focus on randomly mating populations to explore the effect of background selection (BGS) on the linked neutral genetic variation in the changing environment. In particular, we will study the neutral site frequency spectrum (SFS) and neutral heterozygosity.

### 3.1 Multilocus Wright-Fisher model

We use the multilocus Wright-Fisher model to study the effect of recurrent deleterious mutations at the selected site in a randomly mating (*f* = 0), asexual population of size *N*. Consider a large genome in which a single selected site is linked to infinite number of neutral sites. At the linked neutral sites, the mutations occur according to an infinite-sites model, meaning that each mutation occurs at a new site that does not have any mutation before. The population evolves according to discrete-generation Wright-Fisher process (Fisher, 1930; Wright, 1931), described in a following manner. At *t* = 0, we start with a monomorphic population, and subsequent generations are obtained by first reproduction and then mutation. In reproduction, the offspring generation is generated by sampling individuals from the parent generation with equal probabilities. The neutral mutations occur in the population with a rate *u*_*n*_ at a monomorphic site. After a drift-neutral mutation equilibrium for the SFS has been reached, the deleterious mutations are introduced, and the chosen individual acquires deleterious mutations according to the Poisson distribution with mean *Nu*_*d*_. The offspring generation is then generated by picking individuals from the parent’s generation with probabilities proportional to their relative fitnesses. The absolute fitness for an individual with *k* deleterious mutations is given by (1 − *s*(*t*))^*k*^. Unlike in the previous section, here we study the discrete-time model, and the selection changes periodically in the square waveform given by

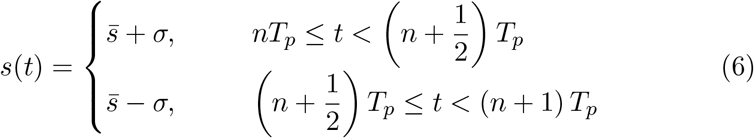

where 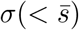 is the amplitude, *T*_*p*_ is the time period of the oscillation and *n* is a positive integer.

To quantify the effect of background selection on the linked neutral diversity, we mainly focus on the two summary statistics, time-averaged site frequency spectrum (SFS), and time-averaged heterozygosity, 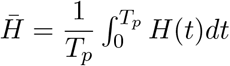. It has previously been shown (Charlesworth, 1993) that in static environment, for strongly deleterious mutations, the effect of the BGS on SFS is same as the neutral expectation with an effective population size, 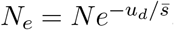, where 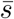 is the magnitude of the selection strength for the deleterious mutation. But for moderately strong selection, the background selection distorts the shape of the SFS from the neutral expectation of monotonically decaying function to a U-shaped function(CvijoviĆ *et al*., 2018). Here we investigate these results in changing environment.

### 3.2 Fluctuating environment between deleterious mutations

We consider the case when the selection coefficient changes between two negative selection coefficients so that the mutation always remains deleterious. We find that, in the slowly changing environment, the time-averaged SFS 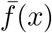 and time-averaged heterozygosity 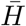 are significantly different compared to the corresponding quantities in the static environment with the time-averaged selection coefficient as depicted in Fig. 3.

**Figure 3:**
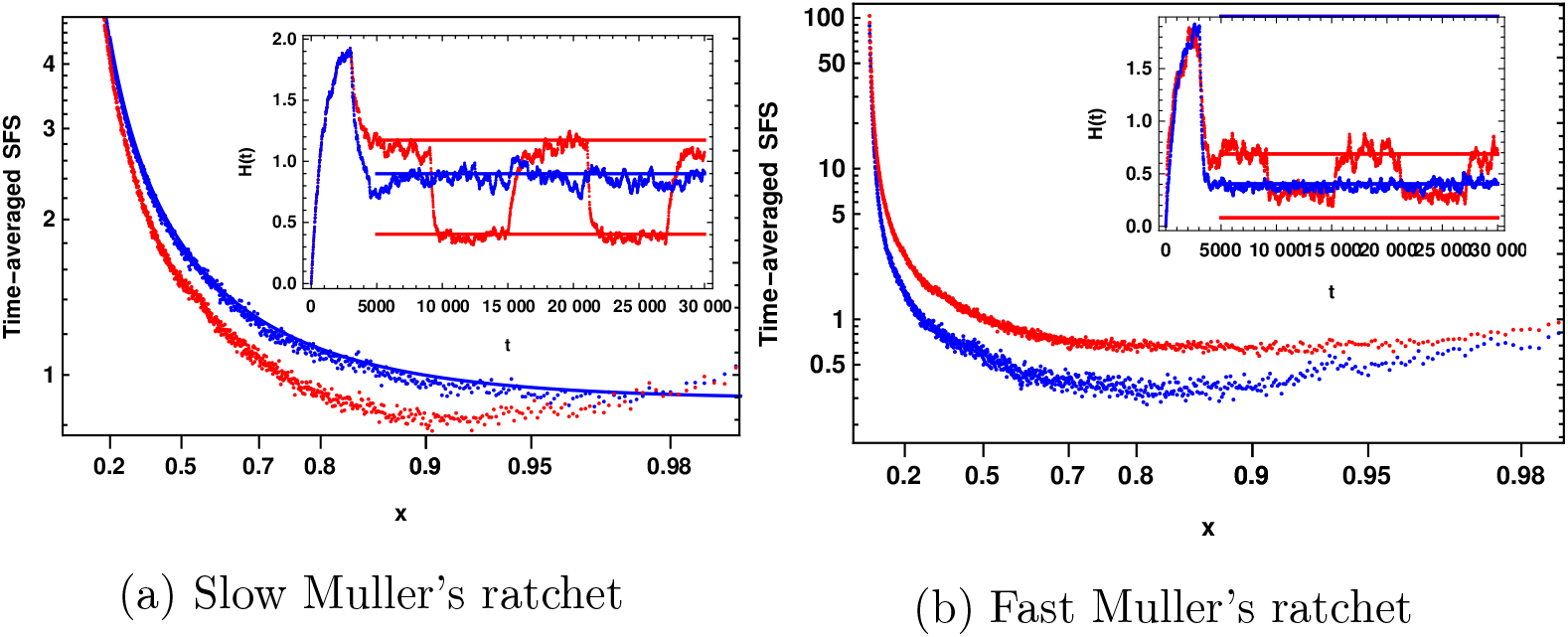
Strictly deleterious mutations: The time-averaged site frequency spectrum for the static environment (blue) and for the slowly changing environment (red) when the selection coefficient is always negative and varies periodically with time period *T*_*p*_ between 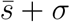 in first half of the cycle and 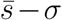 in the second half of the cycle. The parameters are *N* = 1000, *Nu*_*n*_ = 1, *u*_*d*_ = 0.08, 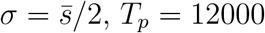, (a) 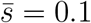 and (b) 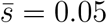. The solid blue line is 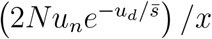. The inset shows the time-dependent heterozygosity for the slowly changing environment (red) and static environment (blue), the solid lines represent heterozygosity for the neutral population with the corresponding effective population sizes. The population evolves under drift neutral mutations equilibrium for SFS till *t* = 3000, and after that, deleterious mutations are introduced.

Figs. 3a, and 3b show respectively that when Muller’s ratchet clicks slowly and rapidly, the time-averaged SFS is smaller and larger, respectively, in the slowly changing environment than the SFS with the time-averaged selection coefficient 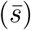 in the static environment. When the neutral allele frequency is very low, the effect of deleterious mutations is negligible, and the time-averaged SFS in changing environment is the same as the neutral SFS, which is analogous to the effect of background selection on neutral SFS in static environment at low frequency (CvijoviĆ *et al*., 2018). However, when the frequency of linked neutral mutations is intermediate, the deleterious mutations are exposed to a changing environment for a longer time, and the SFS at linked neutral sites changes more as a result (Figs. 3a, 3b). In a slowly changing environment, the time-dependent heterozygosity, *H*(*t*), oscillates between the static heterozygosities, 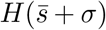 and 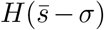, reaching equilibrium in both positive and negative cycles of selection coefficient (inset of Fig. 3).

The effect of changing environment on mean heterozygosity for slow and fast Muller’s ratchet is the same as that on SFS described above. For the slow Muller’s ratchet, 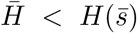, whereas when Muller’s ratchet clicks fast, 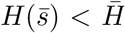. The effect of changing environment is strong for the moderately strong selection 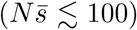 as depicted by Fig. 4, where even in the slowly changing environment, the mean heterozygosity has deviated to 30 − 40% (inset of Fig. 4) from the corresponding heterozygosity in the static environment.

**Figure 4:**
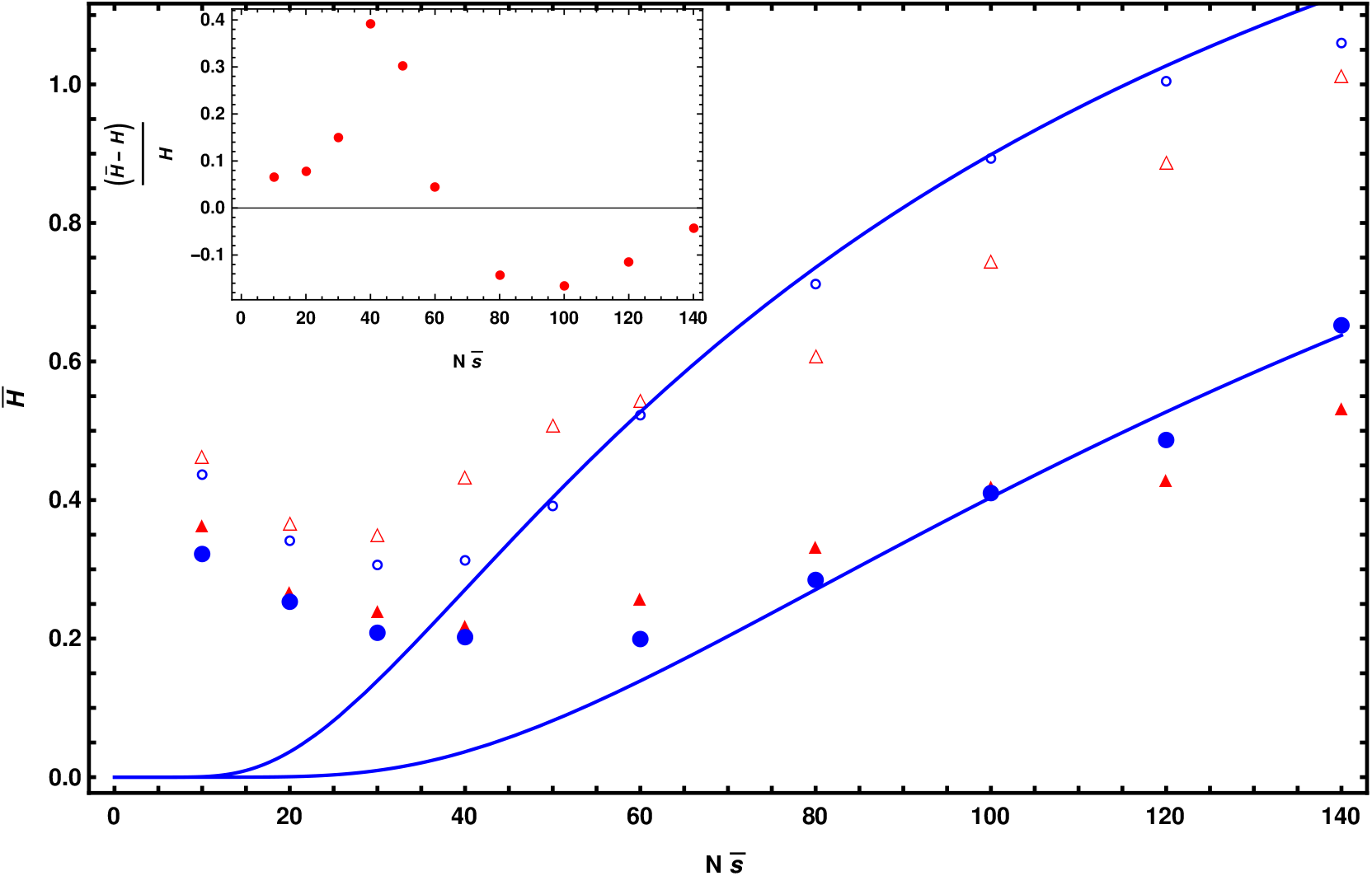
Strictly deleterious mutations: The mean heterozygosity as a function of scaled selection coefficient for static environment (circles) and slowly changing environment with selection coefficient (triangles) varies between 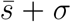 and 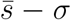 with time period *T*_*p*_. The other parameters are *N* = 1000 (open symbols), *N* = 2000 (closed symbols), *Nu*_*n*_ = 1, *u*_*d*_ = 0.08, 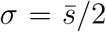 and *T*_*p*_ = 12000 respectively. The solid lines represent mean heterozygosity for the neutral population with effective population size as 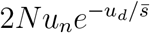. The inset shows the relative change in the mean heterozygosity due to the changing environment as compared to the static environment for *N* = 1000.

When the selection strength is strong 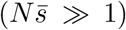, and Muller’s ratchet operates slowly, the time-averaged heterozygosity in the changing environment can be approximated by the mean of heterozygosity in the static environment with selection coefficient 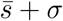, and 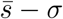,

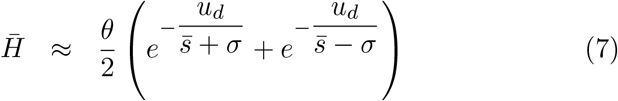

where *θ* is the scaled mutation rate, *θ* = 2*Nu*_*n*_. In the strong selection limit 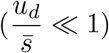, on expanding the *R*.*H*.*S* of above equation, we find

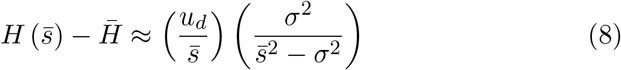

which shows that mean heterozygosity (7) in changing environment is always smaller than that in static environment 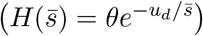, as shown in Fig. 4, and the deviation is given by,

Using the same argument, the time-averaged SFS in the slowly changing environment is smaller than that in the static environment for strong selection, as depicted in Fig. 3a.

When the selection strength is moderately strong, and *σ* is of the order 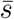 (but smaller than 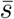), the selection coefficient in the second half of the cycle becomes weak 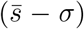, where Muller’s ratchet operates fast, and the mutation classes other than the least loaded class can contribute to the SFS. The mean heterozygosity in fast ratchet regime when 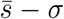 becomes very weak can be approximated by the neutral heterozygosity *θ* whereas the selection in the first half of the cycle, 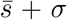, is strong and only the least loaded class contributes to the heterozygosity. The mean heterozygosity is then given by

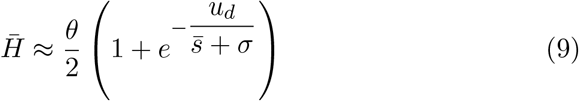

which is greater than the static heterozygosity 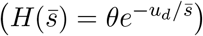.

The time-averaged heterozygosity behaves non-monotonically with selection strength in the changing environment as observed in Fig. 4. The similar dependence is shown by Gordo *et al*. (2002), and Good *et al*. (2014) for the mean number of pairwise differences in the static environment. Since the two regimes of Muller’s ratchet are separated by 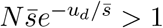 (Gordo and Charlesworth, 2000a; Jain, 2008), the minimum of the mean heterozygosity scales linearly with 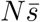 for the fixed 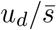 as shown in Fig. 4.

### 3.3 Fluctuating environment between neutral and deleterious mutations

Here, we examine the scenario where the environment changes in such a way that the mutation is deleterious in the first half of the cycle where background selection operates, but neutral in the second half of the cycle where the neutral mutation can get eliminated or fixed due to genetic drift.

Figure 5 shows that, for the moderate selection, the time-averaged SFS in changing environment at the low frequencies (0.01 − 0.05) has a different shape than that in the static environment with a time-averaged selection coefficient, and it is more skewed in the latter case. The contribution of the neutral cycle (*θ*/*x*) to the SFS is larger than that of the deleterious cycle 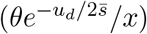, resulting in the time-averaged SFS in the changing environment to have the similar shape to that in the neutral environment as depicted in Fig. 5.

**Figure 5:**
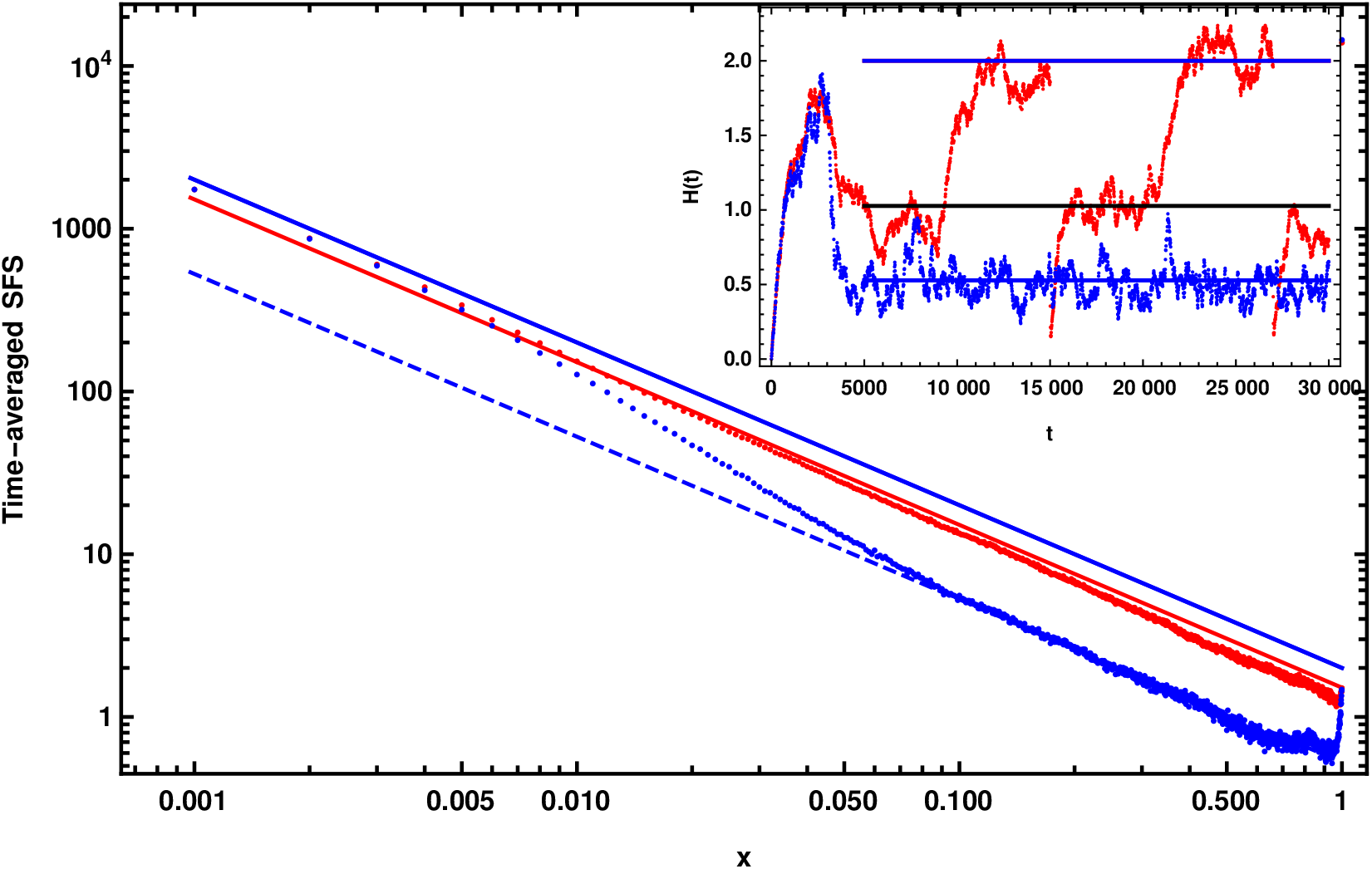
Deleterious and neutral mutations: The time-averaged SFS, when the environment is static (blue) with selection coefficient 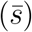 and changes slowly (red) with selection coefficient varies between 0 (neutral) and 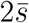 (deleterious), and the Muller’s ratchet clicks slowly in the deleterious cycle. The blue solid and blue dashed line represents SFS for the neutral mutation with population size *N* and 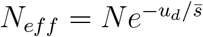 respectively. The red solid line represents the time-averaged SFS given as, 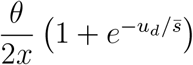. The other parameters are *N* = 1000, *u*_*d*_ = 0.08, *u*_*n*_ = 0.001, 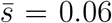, and *θ* = 2*Nu*_*n*_. The inset shows the time-dependent heterozygosity in the slowly changing environment (red) which oscillates between the corresponding heterozygosity in the neutral population with size *N* and *N*_*eff*_ respectively. For the static environment (blue), the heterozygosity in the equilibrium population is given by 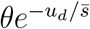. The solid lines from top to bottom represents 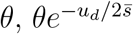 and 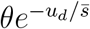 respectively. The population evolves under drift neutral mutations equilibrium for SFS till *t* = 3000, and after that, deleterious mutations are introduced.

Fig. 6 shows that 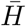 in a slowly changing environment is significantly different from heterozygosity in a static environment 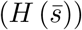 when the selection is moderately strong,

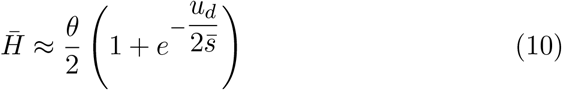

whereas in the static environment, 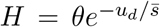 where *θ* = 2*Nu*_*n*_. The mean heterozygosity in slowly changing environment, 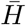, is always greater than that in the static environment, 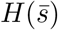, as depicted in Fig. 6. The deviation is approximately (3/4)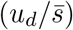 when 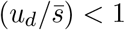. In the slowly changing environment, 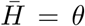 in limits 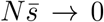 and 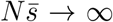 as shown in Fig. 4. The time-averaged heterozygosity follows the non-monotonic behavior with selection strength similar to the previous case where the mutation is always deleterious, and here also, the minimum of the time-averaged heterozygosity scales linearly with 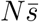 as shown in the inset of Fig. 6. Hence, these results suggest that the levels of genetic variation at the linked neutral sites are significantly different in the changing selective environment for the deleterious mutation than in the static environment.

**Figure 6:**
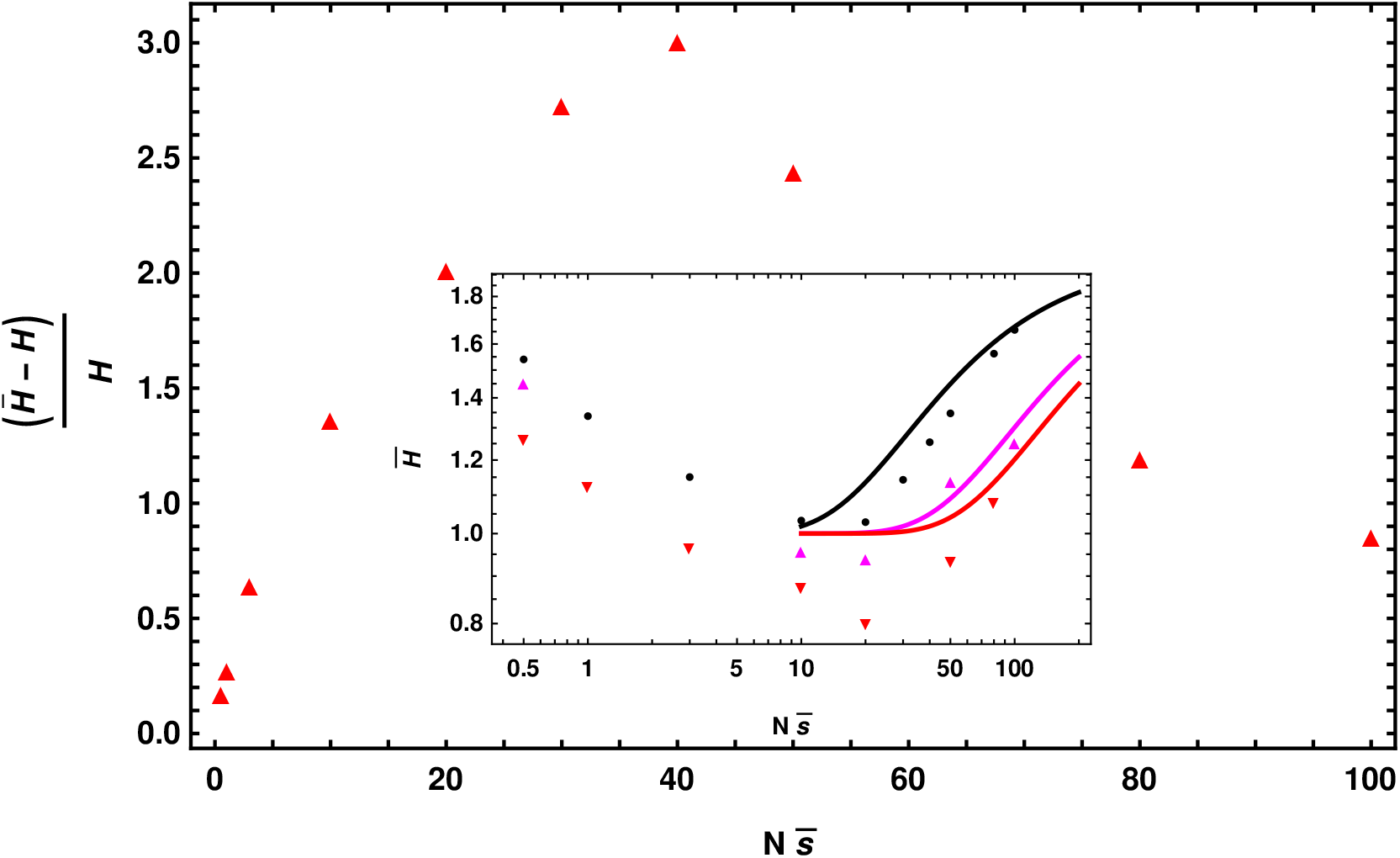
Deleterious and neutral mutations: The relative change in the mean heterozygosity in the changing environment 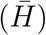 from the static environment (*H*) when selection coefficient varies periodically between zero (neutral) and 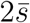 (deleterious) with time period *T*_*p*_. The other parameters are *N* = 1000, *Nu*_*n*_ = 1, *u*_*d*_ = 0.8 and *T*_*p*_ = 12000 respectively. The inset shows that the time-averaged heterozygosity behaves non-monotonically with the scaled selection coefficient and the minimum of the mean heterozygosity scales linearly with 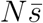. The other parameters are *N* = 1000 (•), *N* = 1500 (▲) and *N* = 2000 (▼) respectively. The solid lines represents 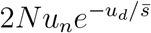 for *N* = 1000, 1500, and 2000 from top to bottom.

## 4 Discussion

In a recent study by Kaushik and Jain (2021), it was found that the Maruyama-Kimura symmetry (Maruyama, 1974; Maruyama and Kimura, 1974) for the conditional mean fixation time in constant environment does not hold in changing environment. In this paper, we have explored this symmetry for the more general case when inbreeding occurs in the population. We find that the deviation in the conditional mean fixation time for both beneficial as well as deleterious sweeps is maximum in the case of randomly mating population. When the inbreeding coefficient is high, the symmetry between beneficial and deleterious conditional mean fixation time breaks very mildly, which makes it difficult to distinguish the diversity patterns generated due to beneficial and deleterious sweeps.

Since the effects of changing environment are more significant in deleterious mutations in the randomly mating population, we investigated the effect of changing environment when recurrent deleterious mutations arise at the selected site and are eliminated from the population (Charlesworth, 1993). When the selection strength is moderate, the effect of background selection on time-averaged SFS is found to be qualitatively different from that in the static environment (CvijoviĆ *et al*., 2018).

When the population fluctuates between zero and a negative selection coefficient in a slowly changing environment, the time-averaged SFS and mean heterozygosity are larger than in a static environment with a time-averaged selection coefficient. Whereas, when the selection coefficient fluctuates in such a way that it is always negative, the time-averaged SFS and mean heterozygosity are smaller (larger) than that in the static environment with a time-averaged selection coefficient when the Muller’s ratchet clicks fast (slow). These findings imply that the recurrent deleterious mutations at the selected site result in qualitatively different levels of genetic variation in the changing environment.

Here, we have studied the effect of changing environment on the SFS due to the background selection using numerical simulations. However, there is a scope to develop the analytical theory to study the behavior of non-equilibrium SFS in the changing environment when Muller’s ratchet operates fast. Here, we have the selected site being either deleterious or neutral; it will be interesting to look at the case where the selected site is beneficial where selective sweeps also occur. Throughout this article, we have assumed that population size is constant, and it will be interesting to explore the effects of demography. Studying the background selection in the changing environment for the selfing population is also interesting because inbreeding is expected to reduce genetic diversity significantly.

## Acknowledgments

S.K. thank his thesis supervisor Kavita Jain for valuable discussions on the work, helpful suggestions and comments on the manuscript. S.K. also thank Himanshu Joshi, Lakshita Jindal, and Sayantan Maity for the comments on the manuscript.

## Appendix

### Appendix A Diffusion Theory

The eventual fixation probability, *P*_f_(*x, t*_0_), that the mutation allele with initial frequency *x* at time *t*_0_ fixes in the population obeys the following backward Kolmogorov equation, and depends upon on the arrival time of the mutation (*t*_0_) (Ewens, 2004; Uecker and Hermisson, 2011; Devi and Jain, 2020) as

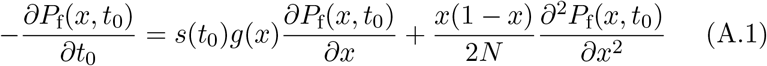

When the mutation is on-average neutral, and environmental change is slow, fixation probability is approximated using perturbation theory as *P*_f_ ≈ *P*_0_ + *NωP*_1_. The *P*_0_ and *P*_1_ are, respectively, the eventual fixation probability in the static environment and the leading order deviation in *ω* in the fixation probability, which obey the following partial differential equations (Devi and Jain, 2020),

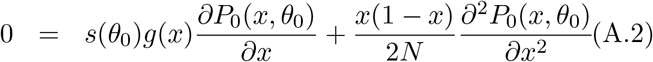

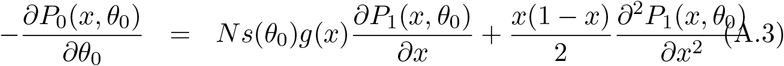

with the boundary conditions as *P*_0_(0, *θ*_0_) = 0, *P*_0_(1, *θ*_0_) = 1, *P*_1_(0, *θ*_0_) = 0, and *P*_1_(1, *θ*_0_) = 0. The *θ*_0_ = *ωt*_0_ is the initial phase at which mutation arises in the population. Equation (A.3) is similar to Eqn. (B.3) in (Devi and Jain, 2020) and can be solved using the integrating factor method for the arbitrary inbreeding coefficient. Using (A.2), the fixation probability in static environment is given by

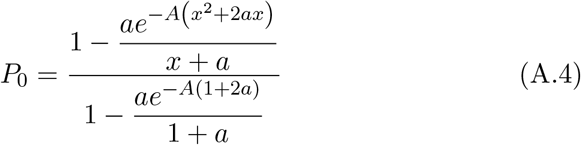

where

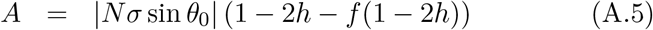

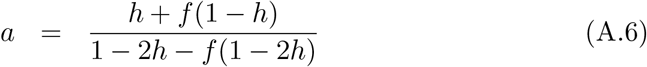

For strong selection |*Nσ*| ≫ 1, and the initial mutation frequency *x* → 0, the *P*_0_, and *P*_1_ are given as

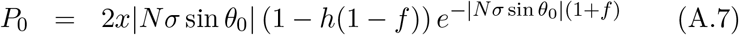

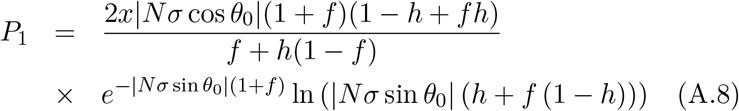

Unlike in a randomly mating population, the deviation in fixation probability is weakly dependent on the dominance parameter when the inbreeding coefficient is high. Since conditional mean fixation time depends upon the fixation probability as discussed in the main text, the deleterious mutations, be it deleterious or recessive, are weakly affected in the slowly changing environment for highly inbred individuals, which is not the same in randomly mating population.

The time inhomogeneous Kolmogorov backward equation used for randomly mating populations in Kaushik and Jain (2021) can be extended easily for inbreeding populations. We use the same perturbation theory used in fixation probability for the conditional mean fixation time when mutation is on-average neutral, by writing 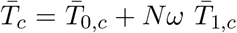 in (1). The 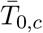 and 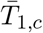 follows (Kaushik and Jain, 2021)

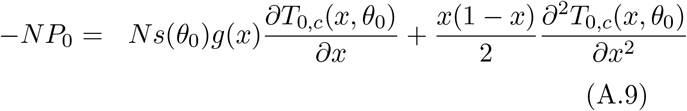

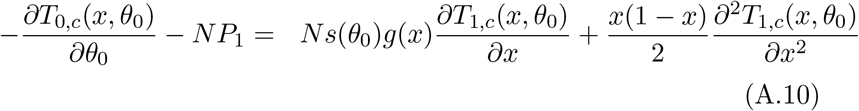

### Appendix B Semi-deterministic approximation

We follow the same procedure as in Martin and Lambert (2015), and Kaushik and Jain (2021) to calculate the fixation time for the beneficial mutations in an inbreeding population. Using (2), the fixation time distribution can be straightforwardly generalized for inbreeding population as

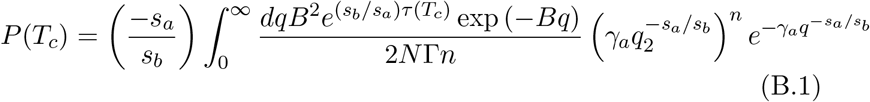

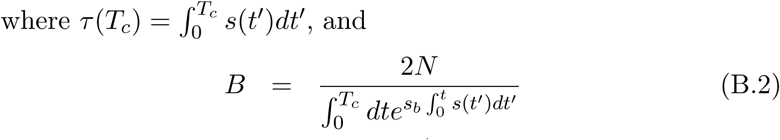

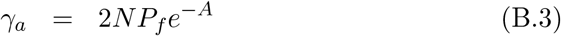

where *P*_*f*_ = *s*_*a*_*σ* sin *θ*_0_ is the eventual fixation probability for a single mutation. For the static environment the fixation time is given by (Martin and Lambert, 2015)

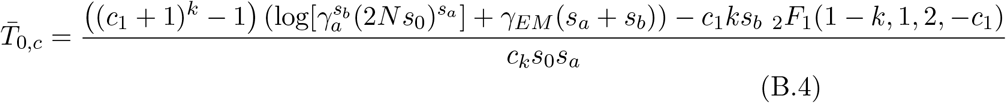

where *s*_0_ = *s*_*b*_*s*(0), *k* and *n* are the initial and the established copies of mutation allele respectively. The _2_*F*_1_(*a, b*; *c*; *z*) is the Gauss hypergeometric function (Abramowitz and Stegun, 1964). The *γ*_*a*_, *c*_1_, and *c*_*k*_ are given by

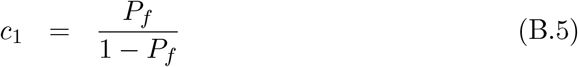

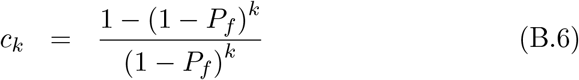

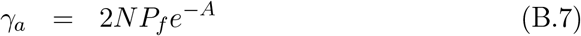

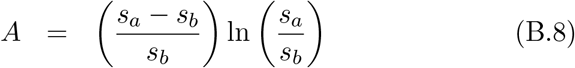

For a single mutation (*k* = 1), the (B.4) matches exactly with Eqn. (17) in GlÉmin (2012) where approximate expression of the conditional mean fixation time is obtained using diffusion theory. The deviation in the conditional mean fixation time in the slowly changing environment, 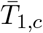 is then given by,

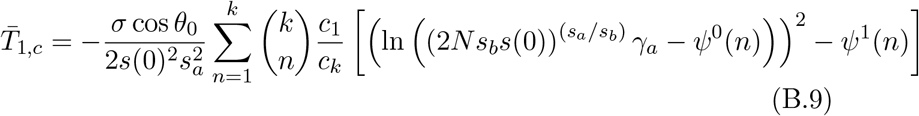

where, *ψ*^*n*^ (*x*) is the Polygamma function, given by the (*n*+1)st derivative of the logarithm of the gamma function (Γ(*x*)) (Abramowitz and Stegun, 1964).

## References

Abramowitz, M. and I. A. Stegun, 1964 Handbook of Mathematical Functions with Formulas, Graphs, and Mathematical Tables. Dover.

Barton, N. H., 2000 Genetic hitchhiking. Philos. Trans. R. Soc. Lond. B 355: 1553–62.

Berry, A. J., J. W. Ajioka and M. Kreitman, 1991 Lack of polymorphism on the Drosophila fourth chromosome resulting from selection. Genetics 129: 1111–1117.

Braverman, J. M., R. R. Hudson, N. L. Kaplan, C. H. Langley, and W. Stephan, 1995 The hitchhiking effect on the site frequency spectrum of DNA polymorphisms. Genetics 140: 783–796.

Buffalo, V., 2021 Quantifying the relationship between genetic diversity and population size suggests natural selection cannot explain Lewontin’s paradox. eLife 10: e67509.

Charlesworth, B. and D. Charlesworth, 2010 Elements of Evolutionary Genetics. Roberts and Company, Colorado.

Charlesworth, B. and J. D. Jensen, 2022 How can we resolve Lewontin’s paradox? Gen. Biol. and Evol. 14: evac096.

Charlesworth, B., M. T. Morgan, and D. Charlesworth, 1993 The effect of deleterious mutations on neutral molecular variation. Genetics 134: 1289–1303.

Cvijović, I., B. H. Good and M. M. Desai, 2018 The effect of strong purifying selection on genetic diversity. Genetics 209: 1235–1278.

Devi, A. and K. Jain, 2020 The impact of dominance on adaptation in changing environments. Genetics 216: 227–240.

Ewens, W., 2004 Mathematical Population Genetics. Springer, Berlin.

Ewing, G., J. Hermisson, P. Pfaffelhuber, and J. Rudolf, 2011 Selective sweeps for recessive alleles and for other modes of dominance. J. Math. Biol. 63: 399–431.

Feller, W., 1951a Diffusion processes in genetics. In Proceedings of the Second Berkeley Symposium on Mathematical Statistics and Probability, pp. 227–246.

Felsenstein, J., 1974 The evolutionary advantage of recombination. Genetics 78: 737–756.

Fisher, R. A., 1930 The genetical theory of natural selection. Oxford: Clarendon Press.

Glémin, S., 2012 Extinction and fixation times with dominance and inbreeding. Theo. Pop. Biol. 81: 310–316.

Good, B. H., A. M. Walczak, R. A. Neher, and M. M. Desai, 2014 Genetic giversity in the interference selection limit. PloS Genetics 10: e1004222.

Gordo, I., A. Navarro, and B. Charlesworth, 2002 Muller’s ratchet and the pattern of variation at a neutral locus. Genetics 161: 835–848.

Gordo, I. and B. Charlesworth, 2000a The degeneration of asexual haploid populations and the speed of Muller’s ratchet. Genetics 154: 1379–1387.

Haldane, J. B. S., 1927 A mathematical theory of natural and artificial selection. V. Proc. Camb. Philos. Soc. 23: 838–844.

Hudson, R. R. and N. L. Kaplan, 1995 The coalescent process and background selection. Phil. Trans.: R. Soc. Lond.B 349: 19–23.

Jain, K., 2008 Loss of least-loaded class in asexual populations due to drift and epistasis. Genetics 179: 2125–2134.

Kaushik, S. and K. Jain, 2021 Time to fixation in changing environments. Genetics 219: iyab 148.

Crow, J. F. and M. Kimura, 2009 An introduction to population genetics theory. The Blackburn Press, New Jersey.

Kimura, M., 1971 Theoretical foundations of population genetics at the molecular level. Theo. Pop. Biol. 2: 174–208.

Lewontin R. C., 1974 The Genetic Basis of Evolutionary Change.. New York: Columbia University Press.

Martin, G. and A. Lambert, 2015 A simple, semi-deterministic approximation to the distribution of selective sweeps in large populations. Theo. Pop. Biol. 101: 40–46.

Maruyama, T., 1974 The age of an allele in a finite population. Genetics. Res. Camb. 23: 137–143.

Maruyama, T. and M. Kimura 1974 A note on the speed of gene frequency changes in reverse directions in a finite population. Evolution 28: 161–163.

Maynard Smith, J. and J. Haigh, 1974 Hitchhiking effect of a favourable gene. Genet. Res. 23: 23–35.

Muller, H. J., 1964 The relation of recombination to mutational advance. Mutat. Res. 1: 2–9.

Risken, H., 1996 The Fokker Planck equation. Methods of solution and applications. Springer, Berlin.

Stephan, W., T. H. E. Wiehe, and M. W. Lenz, 1992 The effect of strongly selected substitutions on neutral polymorphism: analytical results based on diffusion theory. Theor. Pop. Bio. 41: 237–254.

Stephan, W., 2019 Selective Sweeps. Genetics 211: 5–13.

Uecker, H. and J. Hermisson, 2011 On the fixation process of a beneficial mutation in a variable environment. Genetics 188: 915–930.

van Herwaarden, O. A and N. J. van der Wak, 2002 Extinction time and age of an allele in a large finite population. Theo. Pop. Biol. 61: 311–318.

Wright, S., 1931 Evolution in mendelian populations. Genetics 16: 97–159.

